# Mutation of a stromal C-terminal threonine residue of *Photosystem II subunit S* slows down NPQ induction and speeds up relaxation

**DOI:** 10.1101/2025.05.23.655821

**Authors:** Weng Yik Chin, Julia Walter, Alice K. J. Robijns, Johannes Kromdijk

## Abstract

In order to prevent damage by excess light, photosystem antennae can switch to an energy dissipative mode (termed non-photochemical quenching, NPQ). In higher plants, this switch is facilitated by the presence of Photosystem II subunit S (PsbS) protein, which was discovered 25 years ago. While the role of PsbS in induction of NPQ was soon found to require protonation of key glutamate residues facing the thylakoid lumen, a complete understanding of how NPQ is subsequently initiated is still lacking. Recent work on Norway spruce suggests that reversible phosphorylation at a few key residues of PsbS may coincide with the induction of a sustained dissipative state. Here we used *Arabidopsis thaliana* mutant lines to assess the functional implications of phosphorylation at threonine-259, one of the implicated phosphorylatable residues. Using a set of residue changes expressed in the background of PsbS knock-out mutant, *npq4*, we show that neither phosphomimetic, phosphosubstitution, nor phosphonull complementations could rescue NPQ activity to the level of the unperturbed protein. Instead, all residue substitutions at threonine-259 gave rise to significantly impaired induction and accelerated NPQ recovery, while protein accumulation and thylakoid membrane localisation were not affected. We suggest that these results are consistent with a role for the C-terminus in the propensity or stability of hydrophobic interactions between PsbS monomers to form homodimers, or between PsbS and other LHCII proteins to initiate the quenched state.

**Highlight:** Reversible phosphorylation of threonine-259 on PsbS has previously been suggested to play a role in sustained NPQ during cold-acclimation in evergreen conifers. Here we show that both phosphomimetic and phosphonull substitutions at T259 lead to significantly impaired NPQ induction and faster recovery, which may be explained by their impact on hydrophobic interactions between PsbS and other LHCII proteins.

## Introduction

As sessile organisms, plants are constantly exposed to various abiotic stresses such as excess incident light. Over-excitation of plant pigments may lead to the formation of reactive oxygen species, which in turn cause photo-oxidative damage. To mitigate photodamage, the efficiency of the photosynthetic light-harvesting antennae need to be regulated in response to the prevailing light environment, via feedback mechanisms that dissipate excess energy as heat, collectively termed non-photochemical quenching (NPQ; Ruban, 2016). NPQ plays a crucial protective role in plants by fine-tuning the equilibrium between light energy utilisation and dissipation, thereby sustaining photosynthetic efficiency under fluctuating light conditions.

NPQ components can be differentiated based on their induction and relaxation kinetics as well as molecular players involved, including energy-dependent quenching (qE), zeaxanthin dependent quenching (qZ), photoinhibitory quenching (qI), and plastid-lipocalin dependent quenching (qH) (Nilkens *et al*., 2010; Malnoë, 2018; Amstutz *et al*., 2020). Among these NPQ pathways, qE has been most extensively studied, due to its role as the dominant quenching mechanism under dynamic light conditions in higher plants (Niyogi *et al*., 2005). qE is the most rapid NPQ component, responding on a time scale of seconds to minutes in response to fluctuations in light intensity (Zaks *et al*., 2012). Induction of qE requires acidification of the thylakoid lumen, which occurs due to the proton gradient build-up during electron transport. This leads to the protonation of both *Photosystem II subunit S* (PsbS) and *violaxanthin de-epoxidase* (VDE) (Li *et al*., 2004; Fufezan *et al*., 2012). Protonation of VDE activates the xanthophyll cycle, i.e., the reversible conversion of violaxanthin to antheraxanthin and subsequently zeaxanthin, which in turn is suggested to play a role in the quenching of excess energy via qE or qZ (Nilkens *et al*., 2010). In addition to zeaxanthin, PsbS is necessary to induce the full qE response. PsbS belongs to the light-harvesting complex (LHC) family. In contrast to most other LHC proteins with three transmembrane helices, PsbS has four transmembrane helices (**Fig. 1**), flanked by two β-strands on the stromal side, β1 and β2 at the N-terminus and C-terminus, respectively, along with two lumen-exposed amphiphilic helices, H1 and H2. Recent work also pinpointed a further amphipathic 3_10_ short helix domain, H3, which forms from an unstructured luminal loop during lumen acidification (Liguori *et al*., 2019).

**Fig. 1.**
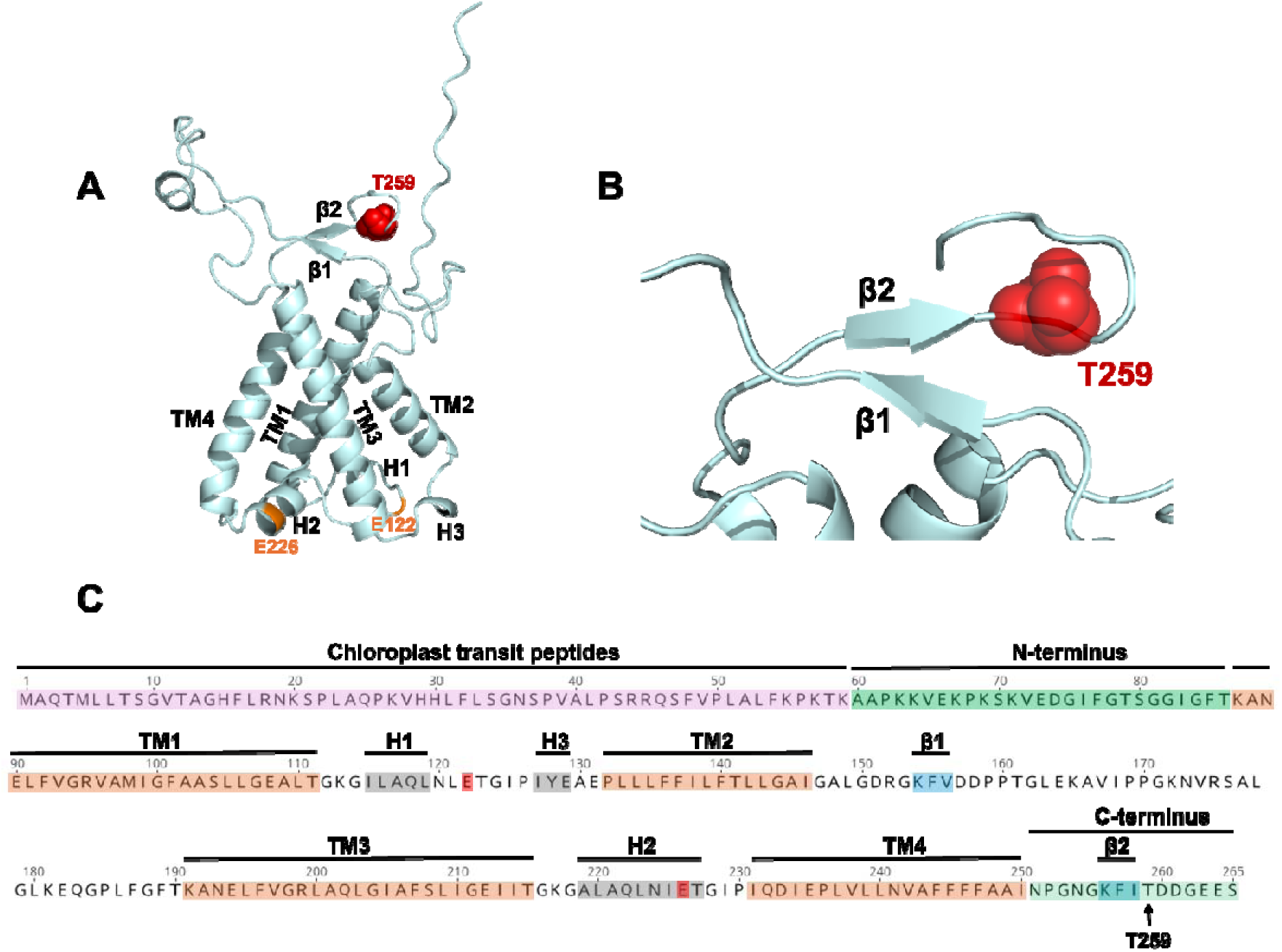
*Arabidopsis thaliana* PsbS structure and sequence. **(A)** Ribbon depiction of the *A. thaliana* PsbS monomer predicted using UniProt: Q9XF91 (without transit peptides) with AlphaFold 3 (Abramson *et al*., 2024). Threonine-259 (T259) residue is highlighted *as* red sphere, whereas the two key pH-sensing glutamate residues, E122 and E226, are highlighted in orange. Transmembrane helices 1-4, helices 1-3, and the two β-strands are denoted as TM1-4, H1-3, and β1-2, respectively. **(B)** Zoomed-in schematic of the β-pleated sheets around the T259 residue. **(C)** Full-length (265 amino acids) annotation of *A. thaliana* PsbS.

Early work indicated that acidification of the thylakoid lumen is sensed through protonation of two glutamate residues (E122 and E226 in *Arabidopsis thaliana*) at the lumen-exposed loops of PsbS (Li *et al*., 2004). E122 lies between H1 and H3, while E226 is positioned within H2. In addition, PsbS undergoes a dimer-to-monomer transition which coincides with qE formation (Bergantino *et al*., 2003), suggesting that the two phenomena may be functionally related (Pawlak *et al*., 2020). Furthermore, molecular dynamics simulations (Liguori *et al*., 2019; Chiariello *et al*., 2023), NMR and NIR spectroscopic observations on recombinant PsbS protein (Krishnan-Schmieden *et al*., 2021) and experimentation with complemented Arabidopsis lines expressing mutated PsbS sequences (Chen *et al*., 2025), have started to elucidate conformational features of the molecular mechanism of PsbS in qE. The consensus view emerging from these studies suggests that at neutral pH, PsbS has higher propensity to form homodimers, while upon lumen acidification, protonation of the key glutamate residues induces conformational changes involving 3_10_ helix formation at the H3 domain and movement of the H2 helical domain from the aqueous to hydrophobic phase, potentially destabilising dimeric PsbS to form monomers. In turn, these changes could favour PsbS interactions with LHCII counterparts, inducing LHCII conformational changes that dissipate excess energy at PSII.

While the regulation of qE by PsbS clearly involves protonation, Grebe *et al*. (2020) recently suggested that PsbS action may also be affected via reversible phosphorylation of several key residues, based on observations in overwintering Norway spruce (*Picea abies*) needles. The reversible phosphorylation of photosynthetic proteins, such as LHCII trimers during state transitions, is well-documented (Rantala *et al*., 2020). However, the work by Grebe *et al*. (2020) is the only work to date suggesting PsbS phosphorylation may have a functional role in NPQ induction. Phosphorylation of PsbS was hypothesised by Grebe *et al*. (2020) to be triggered by acclimation to winter conditions and to aid a sustained quenched state. Evidence came primarily from parallel dephosphorylation and restoration of the unquenched state of PSII antennae, both of which were triggered by acclimation of needles to recovery conditions following seasonal cold exposure. However, further empirical evidence about the functional implications of PsbS phosphorylation are lacking. In addition, while these observations were made in Norway spruce, reversible phosphorylation of PsbS could potentially also occur outside of conifers. Threonine-259 and several other phosphorylated serine/threonine residues described by Grebe *et al*. (2020) are moderately conserved across species. In fact, through phosphoproteomics, Roitinger *et al*. (2015) identified threonine-259 (T259) and serine-265 (S265) as phosphorylatable residues on PsbS in *Arabidopsis thaliana*. However, the role that these residues may play in NPQ has remained untested.

Here, we use a genetic approach to study the impact of PsbS phosphorylation on NPQ induction and relaxation in *Arabidopsis thaliana*. We focused on the T259 residue, which is located close to the stromal C-terminus of PsbS immediately downstream of the β2 strand (**Fig. 1**). Using complementation of the *Arabidopsis npq4-1* mutant background, which lacks PsbS (Li *et al*., 2000), we tested several PsbS sequences with point mutations at T259, in which the threonine residue was substituted to test the impact of structurally mimicking constitutive lack of phosphorylation (alanine, A; phosphonull), substitution with a different phosphorylation site (serine, S; phosphosubstitution), and constitutive phosphorylation (aspartate, D and glutamate, E; phosphomimetics). Surprisingly, in comparison to the control lines, chlorophyll fluorescence measurements showed that all the T259 modified mutants exhibited severely reduced NPQ induction as well as accelerated relaxation, with the phosphomimetic constructs (T259D and T259E) being most strongly impacted. Parallel transcript and protein analyses confirmed similar expression and proper localisation of PsbS within the thylakoid membrane across all lines, suggesting that the NPQ phenotypes of the residue swaps were associated with protein function, although not in a way consistent with regulation via phosphorylation. While previous studies have shown that PsbS capacity to facilitate qE can be significantly impaired by site-directed changes to two key lumen-positioned glutamate residues (Li *et al*., 2000), this is the first time a stromal residue is directly implicated in NPQ kinetics. The profound impact of all residue changes shows that the C-terminus configuration may be important for hydrophobic interactions which could either impact the propensity of the PsbS protein to undergo dimer-to-monomer transitions or the stability of PsbS interactions with PSII antenna proteins.

## Materials and methods

### Plant materials and propagation

Arabidopsis wild-type Columbia-0 (Col-0) and *PsbS-ko* mutant, *npq4-1* (N66021, hereafter *npq4*) seeds were sourced from the Nottingham Arabidopsis Stock Centre (NASC), both were included in this study as reference and negative control, respectively. PsbS point mutations were generated by swapping the T259 residue to alanine, aspartate, glutamate, and serine (hereafter T259A, T259D, T259E, and T259S, respectively) via site-directed PCR mutagenesis, conferring the predicted sequence changes in **Table S1**. All PsbS mutant sequences, as well as the native PsbS coding sequence as a control, were ligated to the *A. thaliana Rubisco small subunit 1A* (RbcS1A) promoter and *heat shock protein 18.2* (HSP) terminator for robust expression into the pL1V-F2 vector via Golden Gate cloning (Weber *et al*., 2011). Level 2 constructs also contained an mRuby3 fluorescence marker module (FAST, Shimada *et al*., 2010) under the control of the *A. thaliana oleosin 1* (OLE1) promoter and *nopaline synthase* (NOS) terminator to enable rapid screening of transformants via constitutive seed-coat expression. All constructs were used to transform *npq4*, which lacks PsbS, via floral dipping. Homozygous mutant lines from the T3 generation were genotyped via PCR (^5’^TCTGGAAACTCTCCGGTTGC^3’^ and ^5’^CCACAAATTCATAACACAACAAGCC^3’^) and confirmed by Sanger sequencing (**Fig. S1**). For experiments detailed below, homozygous seeds from three independent events per construct were sown in Levington Advance Seed & Modular F2 compost and sand with 3:1 ratio, then stratified at 4 °C for 48 hr. After 2 weeks, seedlings were transplanted into individual pots and grown under ∼ 200 µmol photons m^−2^ s^−1^ (8 hr light: 16 hr dark photoperiod) at 22 °C, 60% relative humidity in a controlled growth chamber. Unless otherwise specified, all measurements were performed on young fully expanded leaves from 6-week-old plants.

### Chlorophyll fluorescence imaging

Chlorophyll fluorescence measurements were carried out using a closed chlorophyll fluorescence imaging system (FluorCam FC 800-C, Photon Systems Instruments, Czech Republic). Plants were dark-adapted for 1 hr prior to measurements. One fully expanded young leaf was excised from each biological replicate, placed on wet filter paper, and sandwiched with glass panels. The resulting stacks were dark-adapted for another 15 min inside the fluorescence imaging system, following which, dark-adapted minimal fluorescence yield (Fo) was determined prior to an 800 ms pulse of 4000 μmol photons m^-2^ s^-1^ for maximum fluorescence yield (Fm) measurement. The maximum dark-adapted PSII efficiency was estimated as Fv/Fm = (Fm−Fo)/Fo. Subsequently, NPQ induction kinetics were determined during illumination with actinic white light (1500 μmol photons m^-2^s^-1^) for 600 s, interspersed with 800 ms pulses of 4000 μmol photons m^-2^s-^1^ timed at 20 s, 40 s, 60 s, 120 s, 240 s, 360 s, 480 s, and 600 s. The 600 s illumination was followed by 300 s dark relaxation, in which further 800 ms pulses were timed at 20 s, 40 s, 60 s, 180 s, 300 s.

### Transcript levels and protein expression

One whole leaf per plant was excised, snap-frozen in liquid nitrogen and used for mRNA and protein extraction using the NucleoSpin RNA/protein purification kit (Item No. 740933.50, Macherey-Nagel, GmbH & Co. KG, Düren, Germany) as per manufacturer instructions. RNA concentration of the samples was determined using Nanodrop 2000 spectrometer (ThermoFisher Scientific Inc.), whereas total protein concentration was determined based on the Karlsson method (Karlsson *et al*., 1994). 1000 ng/μl of complemented DNA (cDNA) was synthesised using 500 ng of RNA with the iScript cDNA synthesis kit (Cat no. 1708890, BIO-RAD Laboratories Inc.) according to manufacturer instructions. Real-time quantitative PCR (RT-qPCR) was performed using a reaction mix of 50% (v/v) SsoAdvanced Universal SYBR Green Supermix (Cat no. 1725270, BIO-RAD Laboratories Inc., California, United States), 100 ng cDNA and 250 nM forward/reverse primer. The qPCR reaction was carried out using a real-time thermal cycler (CFX Connect Real-Time System, BIO-RAD Laboratories Inc.) with the amplification protocol of 95 °C 3 min → 39x (95 °C 10 sec → 60 °C 30 sec). ΔCq for each line was calculated based on the difference between the cycle threshold (Cq) values of PsbS and the geometric mean of both reference genes, actin and ubiquitin (for primers, see **Table S3**).

For the extraction of thylakoid membrane (TM) protein samples, whole rosettes of five 10-week-old plants were harvested and pooled per line and snap-frozen prior to extraction. Extraction followed the protocol by Järvi *et al*. (2011) with slight modifications. To minimise sample degradation, extraction steps were performed under dim light and on ice. The final chlorophyll (*chl*) concentration of TM samples was estimated from absorbance at 650.0 nm, 665.0 nm, and 750.0 nm, computed using the following formula: Chlorophyll *a* (μg/ml) = 16.29 x (A_665.2_ - A_750.0_) – 8.54 x (A_650.0_ - A_750.0_). Absorbance was measured with a spectrometer (UV300 UV-Visible Spectrometer, ThermoFisher Scientific Inc., Massachusetts, United States). For protein solubilisation, TM samples were mixed with equal volume of 2x Laemmli sample buffer containing urea (4% (w/v) SDS, 20% (v/v) glycerol, 120 mM Tris-HCl pH 6.8, 6 M Urea, 10% (v/v) β-mercaptoethanol, and 0.05% (w/v) bromophenol blue) at 95 °C for 5 min, then centrifuged with 5000 rcf at room temperature for 5 min before loading onto SDS-gels.

5 μg total protein and 1 μg *chl* TM samples were separated on SDS-PAGE (12% Mini-PROTEAN® TGX™ Precast Gels, BIO-RAD Laboratories Inc.), and blotted onto polyvinylidene difluoride (PVDF) membranes with a rapid transfer system (Trans-Blot Turbo Transfer System, BIO-RAD Laboratories Inc.). Blots were washed in Tris-buffered saline, TBS (88 mM Tris base, 2.5 M NaCl, adjusted to pH 7.5 with HCl), blocked with 5% milk in TBS with 0.05% (v/v) Tween 20 detergent (T-TBS) for one hour, washed twice with T-TBS and incubated overnight with primary antibodies in 1% (w/v) milk in T-TBS shaking at 4°C overnight. The primary antibodies used in this study were anti-PsbS (AS09533, 1:2000 dilution, Agrisera, Sweden) and anti-actin (AS132640, 1:3000 dilution, Agrisera, Sweden). Then, blots were washed four times with T-TBS, incubated in secondary antibody (goat anti-rabbit IgG (H&L) HRP conjugated, 1:10000 dilution, AS09602, Agrisera, Sweden) for one hour and washed three times with T-TBS and twice with TBS. Finally, blots were incubated in the chemi-luminescence substrate (Clarity Western ECL substrate, BIO-RAD) for 5 min prior to imaging. All the imaging was performed with G:Box chemi XRQ gel doc system (Syngene, India). For total protein staining, membranes were incubated with Coomassie stain (0.05% (w/v) Coomassie Brilliant Blue G-250, 50% (v/v) methanol, 10% (v/v) glacial acetic acid) and destained with fixing solution (40% (v/v) methanol, 10% (v/v) acetic acid). To analyse Western blots semi-quantitatively, densitometry using ImageJ was performed as described by Stael *et al*. (2022).

### Statistical analyses

All statistical analyses were carried out using R (Version 4.4.0, R Core Team, 2021) on RStudio (Posit Team, 2024). For “area under curve” calculations of NPQ timeseries during induction (0-600 s, ≤600 s) and relaxation phases (600-900 s, ≥600 s), the “bayestestR” package was employed and data were computed using the “trapezoid” method (Makowski *et al*., 2019). To estimate the NPQ decay rate, NPQ data from the relaxation phase was used to estimate the parameters from the non-linear exponential decay model below using the nls (nonlinear least squares) function in R:

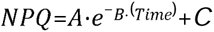

where *A* = amplitude of NPQ decay, *B* = decay rate constant, Time = time-point during relaxation phase, and *C* = residual NPQ. For each parameter, a nested analysis of variance (ANOVA) was performed (**Table S4**), with “line” nested within the “genotype” (construct), to assess differences between genotypes and measured parameters. Significant genotype effects (α = 0.05) were followed up with Tukey’s HSD pairwise multi-comparison test. For densitometry analyses, one-way ANOVA was applied to determine the significance of genotype effects on the basis of grouping individual samples for each line by construct, excluding Col-0. Parameters that showed significant effects of genotype were further examined using Dunnett’s multi-comparison test, with the control construct designated as reference. For PsbS alignment of different species, sequences were obtained from Phytozome v13.0 (https://phytozome-next.jgi.doe.gov/, Goodstein *et al*., 2012), the One Thousand Plant Transcriptomes (1KP) Project (https://db.cngb.org/onekp/, Carpenter *et al*., 2019; One Thousand Plant Transcriptomes Initiative, 2019), and NCBI (https://www.ncbi.nlm.nih.gov/), then aligned using the “ClustalOmega” method from the “Biostrings” package (Pagès *et al*., 2024) and plotted using “ggmsa” (Yu, 2022; Zhou *et al*., 2022).

## Results

### Compromised NPQ induction across T259 mutated lines

In order to assess the impact of phosphorylation at T259 on NPQ induction and relaxation, *npq4* was transformed with T259A (phosphonull), T259S (phosphosubstitution), T259D, T259E (both phosphomimetic), and the native PsbS sequence, all under the strong RbcS1A promoter. NPQ kinetics of three independent homozygous lines per construct were characterised (**Fig. 2**). NPQ induction in the three PsbS control lines was clearly stronger than in Col-0 and *npq4* (**Fig. 2A**), which were included for reference and negative control, respectively, reflecting stronger expression of PsbS driven by the RbcS1A promoter, relative to the native promoter in Col-0. Strikingly, all T259 modified lines exhibited reduced NPQ induction compared to lines expressing the PsbS control sequence (**Fig. 2B-E**), showing markedly slower induction of NPQ. Lines expressing the T259A and T259S constructs showed NPQ levels similar to Col-0 at 600 s, whereas NPQ induction in both phosphomimetic mutants, T259D and T259E, was most strongly impaired, implying a greater impairment of NPQ dynamics arising from phosphomimetic modification. Importantly, consistent Fv/Fm values (**Fig. S2**) of ∼0.83 across all genotypes demonstrated that these differences were not attributable to variation in the dark-adapted values. At 600 s, NPQ of T259A (2.76 ± 0.05) and T259S (2.82 ± 0.04) lines was significantly decreased (Tukey’s HSD: P<0.05) by 43% relative to lines expressing the control PsbS (4.92 ± 0.06) (**Fig. 3A**). At the same time-point, NPQ for both the phosphomimetic lines, T259D (2.00 ± 0.05) and T259E (1.98 ± 0.03) was decreased by 60% relative to lines expressing the control PsbS and was even significantly lower than NPQ in Col-0 (P<0.05) at the same time-point (2.48 ± 0.04). In fact, whereas NPQ induction in Col-0 initially proceeded rapidly and then plateaued, slow induction of NPQ in lines expressing mutations at T259 meant that integrated NPQ throughout the 600 s induction phase was significantly lower in all mutated sequences (P<0.05), not just compared to lines expressing unmodified PsbS, but also compared to Col-0 (**Fig. 3B**). In line with differences in NPQ at 600 s, integrated (area under the curve) values for NPQ were significantly lower in T259D and T259E (Tukey’s HSD: P<0.05), compared to T259A and T259S.

**Fig. 2.**
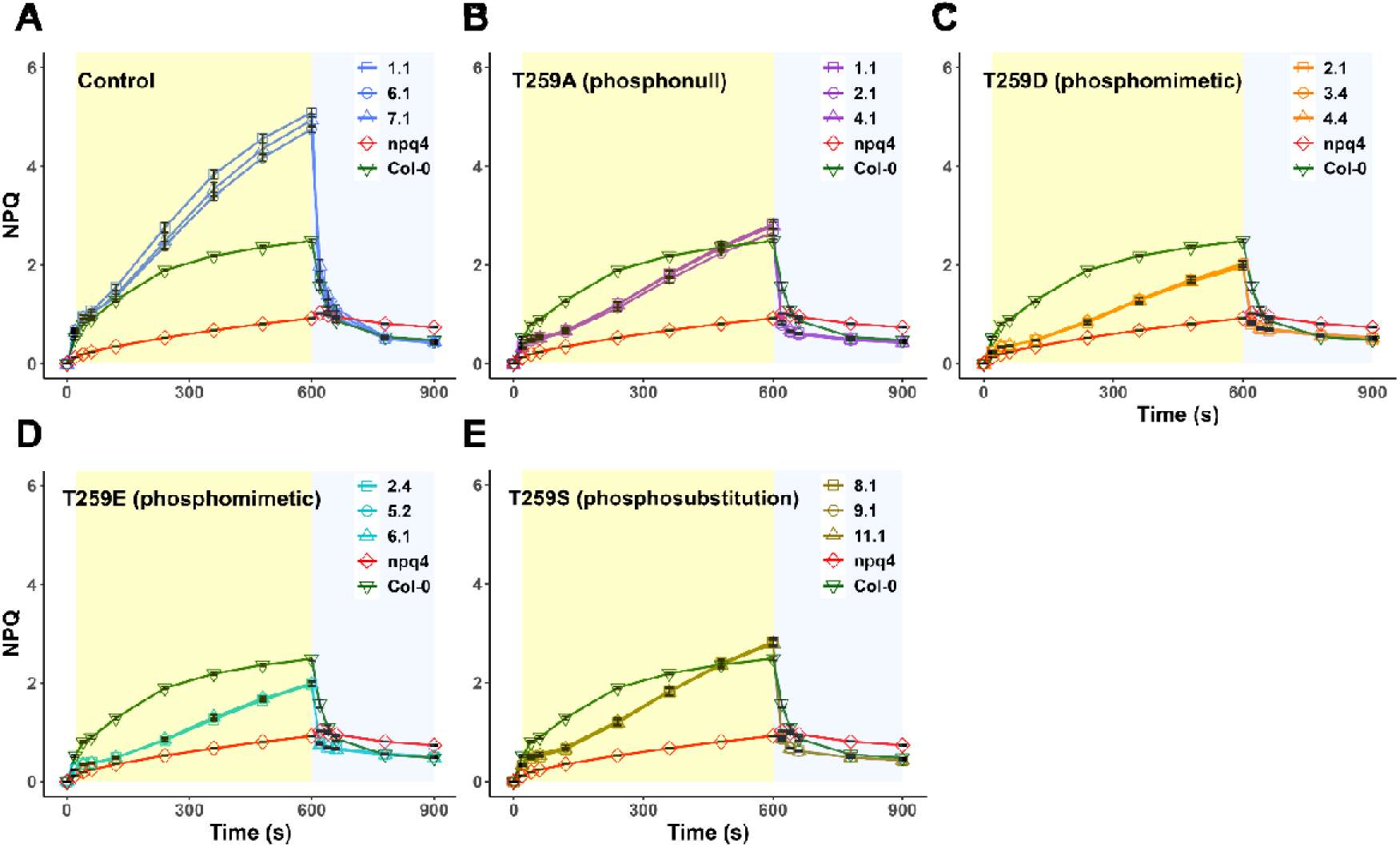
NPQ kinetics of the PsbS mutants. NPQ kinetics of the **(A)** control lines and threonine-259 modified PsbS mutants **(B)** T259A, **(C)** T259D, **(D)** T259E, and **(E)** T259S subjected to 600 s high light induction (yellow shading), followed by 300 s relaxation (light blue shading). Three independent lines are included for each construct. The predicted functional effects of mutants are denoted in parentheses. NPQ of wild-type Col-0 and PsbS-*ko* (*npq4*) mutant are replotted in each panel as references. Each point represents the mean ± SEM of biological replicates (n=8-16) from the respective line.

**Fig. 3.**
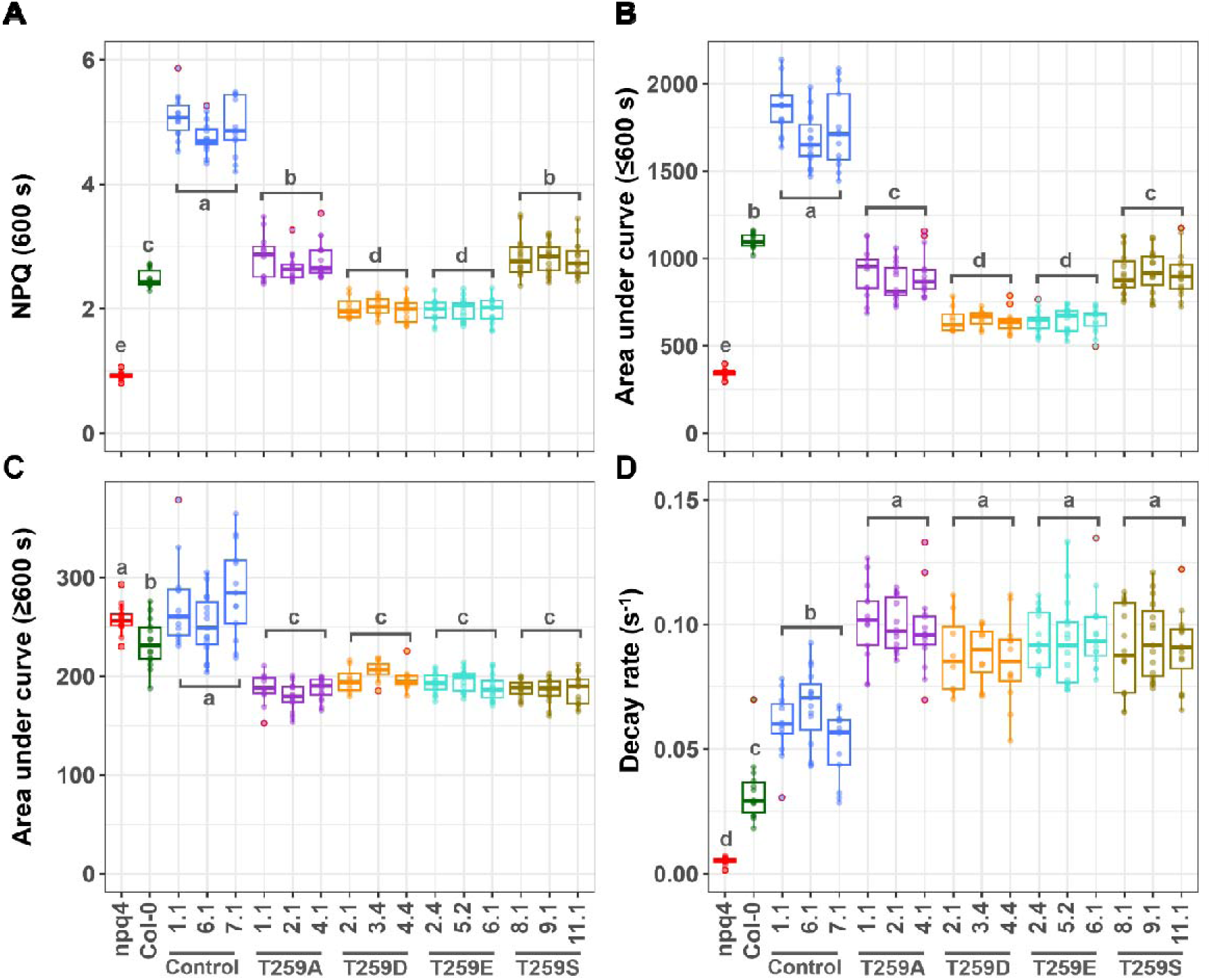
Parameters derived from the NPQ induction and relaxation kinetics of PsbS mutants. **(A)** NPQ value at 600 s, the last measurement pulse during the induction phase. Area under curve determined from the **(B)** induction phase (≤600 s) and **(C)** relaxation phase (≥600 s). **(D)** Decay rates across all genotypes during the relaxation phase estimated via exponential decay models. For all constructs, three independent lines were measured, where n=8-16 biological replications for each line. Outliers are depicted in red circles. Genotypes with the same letter are not significantly different using Tukey’s HSD (α = 0.05) performed on nested ANOVA models, where “line” is nested within “genotype”.

### Accelerated NPQ relaxation in T259 mutated lines

Interestingly, mutation of T259 did not only impair NPQ induction, but also accelerated NPQ relaxation during the final 300 s dark phase. Area under the curve values of NPQ from 600-900 s were significantly lower for all the T259 modified lines (P<0.05) in comparison to lines expressing the control sequence, as well as compared to *npq4* and Col-0 (**Fig. 3C**). This difference occurred despite similar (Col-0) or substantially lower (*npq4*) NPQ values at the start of the dark phase at 600 s (**Fig. 2**), suggesting that the rate of NPQ relaxation was increased in the T259 mutants. Indeed, fitting an exponential decay model to the relaxation phase showed that the NPQ decay rate ranged between 0.085 ± 0.005 s^-1^ to 0.102 ± 0.004 s^-1^ in the mutated T259 lines, which was significantly higher compared to the lines expressing the control PsbS sequence with an average of 0.060 ± 0.002 s^-1^ (P<0.05; **Fig. 3D**), which in turn were significantly higher than Col-0 (0.033 ± 0.003 s^-1^) and *npq4* (0.005 ± 0.0003 s^-1^; P<0.05), probably due to differences in PsbS expression. Hence, regardless of the substituted amino acid, all modifications of T259 significantly increased NPQ relaxation kinetics.

### Elevated PsbS transcript levels and protein abundance across all complemented *npq4* **lines**

To confirm that the differences in NPQ induction and relaxation were directly associated with the T259 mutations and not with quantitative differences in PsbS expression, transcript and protein levels were first assessed in whole leaf extracts. Significantly higher levels of PsbS transcript were found across all control and mutant lines in comparison to Col-0 (P<0.05), which had almost two-fold lower ΔCq values (**Fig. 4A**). However, there was no significant difference between the ΔCq values of control lines and any of the T259 modified lines (Dunnett’s test, P>0.05), suggesting that the varying NPQ kinetics observed in the mutants were not attributable to variation in PsbS transcript levels.

**Fig. 4.**
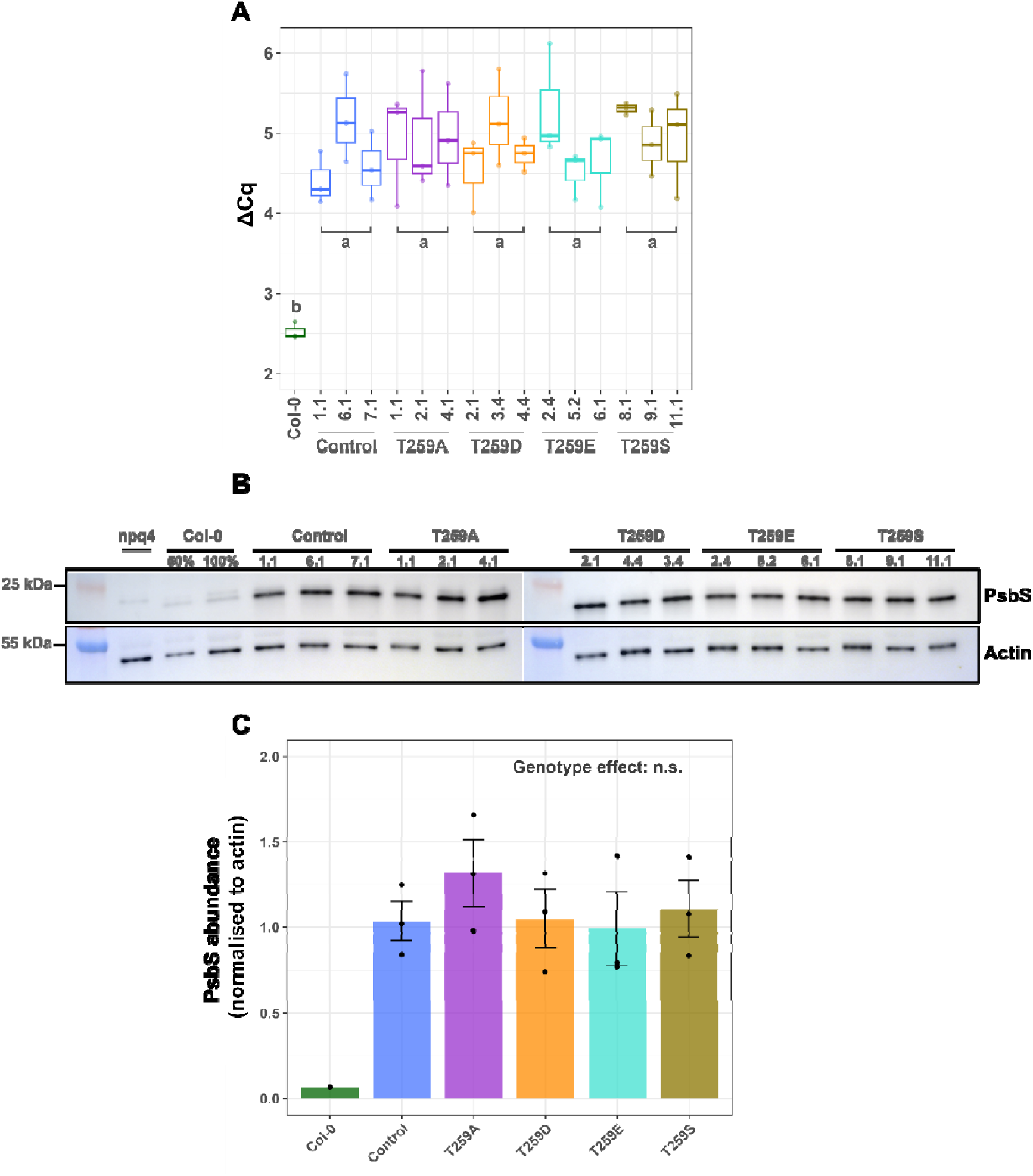
Transcript and total protein western blot analyses. **(A)** ΔCq attained from the difference of RT-qPCR quantification cycle (Cq) values of PsbS relative to the geometric mean of both reference genes, actin and ubiquitin (n=3). Genotypes with the same letter are not significantly different using Tukey’s HSD at α = 0.05 performed on nested ANOVA models, where “line” is nested within “genotype”. **(B)** Western blots against PsbS (∼22 kDa) and actin (∼50 kDa) from total protein extracts of an individual plant from each independent line (n=1). Samples were loaded on the equal basis of 5 μg total protein, loading scales of 50% and 100% Col-0 sample are included in both runs. **(C)** Protein abundance of PsbS based on densitometry analysis from the blotting results. PsbS abundance of respective genotypes is computed by normalising against their corresponding actin abundance. Bars represent mean ± SEM of the three lines for each genotype (n=3), apart from Col-0 (n=1). One-way ANOVA (without Col-0) showed no significant (P>0.1) effect of genotype on PsbS abundance.

Analysis of PsbS abundance in whole-leaf protein extracts showed a similar pattern (**Fig. 4B**). Western blots using a PsbS-specific antibody confirmed that PsbS (∼22 kDa) was expressed strongly in all control and T259 mutant lines (**Fig. 4B**), all of which showed significantly higher PsbS accumulation than Col-0 (P<0.05; **Fig. 4C**). As expected, the PsbS signal was absent in the *npq4* sample, while western blots against actin (∼50 kDa) showed the same band intensity between genotypes (**Fig. 4B**), including *npq4* and Col-0. Quantification via densitometry analysis revealed no significant differences in PsbS protein level between the control lines and T259 mutants (Dunnett’s test, P>0.05) (**Fig. 4C**). Thus, consistent with the analysis of transcript levels, western blot analyses corroborated that the observed NPQ phenotypes described earlier are not associated with accumulation differences in either transcripts or protein of PsbS in the mutant lines.

### Localisation of PsbS in the thylakoid membrane is not affected by mutation of T259

Next, we checked if the modification of T259 may have affected thylakoid localisation. Thylakoid membrane (TM) extracts were separated via SDS-PAGE, blotted onto membrane and probed with the PsbS antibody. Similar to the analyses with whole leaf protein extracts, PsbS abundance in TM extracts was higher in all T259 mutant and control lines relative to Col-0 (**Fig. 5A**). Densitometry analysis of the TM PsbS western blots showed no significant differences (Dunnett’s test, P>0.05) in PsbS abundance between PsbS control lines and the T259 mutant lines (**Fig. 5B**), apart from T259S lines which had significantly higher PsbS abundance than the PsbS control lines (Dunnett’s test, P=0.009). Despite these differences, the observed accumulation of PsbS in the thylakoid membrane across all T259 mutant sequences demonstrates that diminished NPQ induction T259 mutants was not explained by different subcellular localisation patterns of PsbS. These findings therefore strengthen our conclusion that the modifications of T259 impacted PsbS function in induction and relaxation of NPQ, rather than protein accumulation or subcellular localisation.

**Fig. 5.**
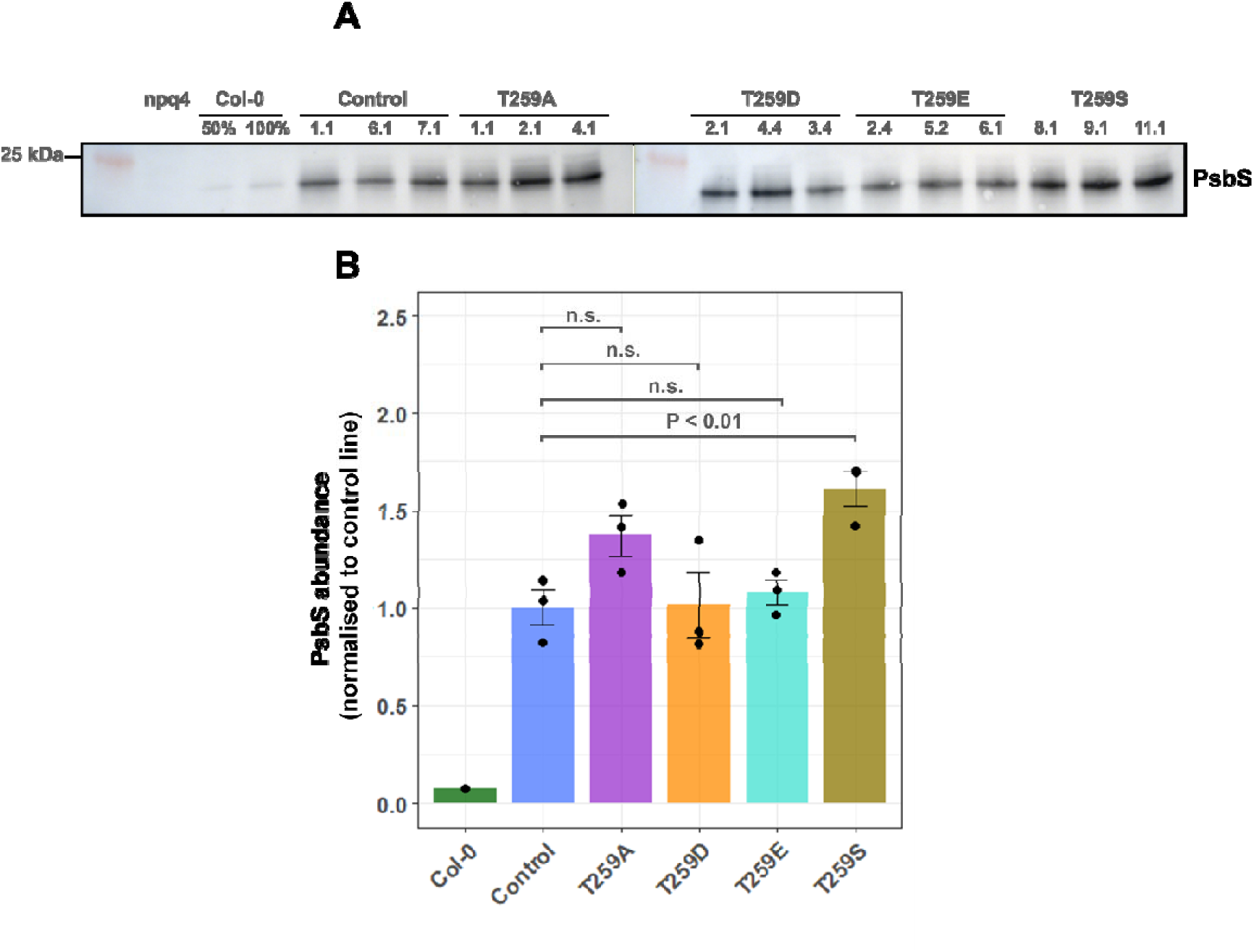
PsbS western blot of thylakoid membrane extracts. **(A)** PsbS western blots performed using the thylakoid membrane extracts pooled from five individual biological replicates of respective lines. Samples are loaded on the equal basis of 1 μg chl, loading scales of 50% and 100% Col-0 sample are included. **(B)** PsbS abundance based on densitometry analysis from the blotting results, all genotypes are normalised against the PsbS abundance of control lines. Bars represent mean ± SEM of three different lines for each genotype (n=3), apart from Col-0 (n=1). One-way ANOVA (without Col-0) showed a significant (P<0.05) effect of genotype on PsbS abundance. P-values above each comparison with the control lines represent significance levels based on Dunnett’s multi-comparison test.

## Discussion

Extensive efforts to understand the precise molecular mechanism of PsbS have been ongoing ever since its crucial role in qE was established. The vast majority of studies have focused on the identification of protonatable residues which may govern conformational responses to changes in lumen pH (Li *et al*., 2004; Liguori *et al*., 2019; Krishnan-Schmieden *et al*., 2021; Chiariello *et al*., 2023). In contrast, here we followed up a putative alternative mode of regulation involving phosphorylation of the stromal residue T259, which was observed by Grebe *et al*. (2020) for PsbS in overwintering needles of Norway spruce. While our mutational analysis showed distinct effects of T259 substitutions on NPQ induction and relaxation kinetics, these did not follow the predicted structural mimicry of (de-)phosphorylated states at this residue. Instead, all T259-mutated PsbS sequences led to diminished NPQ induction and more rapid NPQ relaxation. To our knowledge, this is the first time that such a pronounced effect is observed for a stromal residue, which is likely to provide new leads in unravelling the mechanism of NPQ-facilitation by PsbS.

### NPQ induction is strongly compromised by residue swaps at T259

It is well-known that NPQ amplitude is positively correlated with PsbS abundance (e.g. Li *et al*., 2002 a,b). However, reduced NPQ induction across T259 mutants did not coincide with lower PsbS transcript and protein abundances. On the contrary, all lines showed significantly higher transcript levels and protein expression than Col-0, probably reflecting differences in expression strength between pRbcS1A and the native promoter. Consistent with the observations on whole leaf extracts, TM extracts also showed increased PsbS protein accumulation in all mutated and control T259 lines, relative to Col-0. Altogether these findings convincingly show that the NPQ phenotypes could not be explained by differences in PsbS accumulation or subcellular localisation and instead reflect the functional implications of mutating the T259 residue.

T259A was included in the experiments to represent the non-phosphorylated state, since alanine cannot be phosphorylated. Based on the Grebe *et al*. (2020) findings, this would be predicted to be the NPQ active state, however T259A lines were clearly impaired in NPQ induction. This may imply that there is a structural requirement for threonine at this site, which seems to be confirmed by observations on T259S, which showed a very similar phenotype to T259A. Serine and threonine are both phosphorylatable polar amino acids, but our findings show that these amino acids are not functionally interchangeable at T259. These findings are not uncommon, and threonine and serine are often found to be dissimilar at the molecular level. For example, S891T modification of Brassinosteroid Insensitive 1 (BRI1) did not fully recuperate brassinosteroid signalling in *A. thaliana*, due to slower autophosphorylation at the mutated residue (Oh *et al*. 2015). In addition, structural and functional effects following phosphorylation may also vary between threonine and serine (Ubersax and Ferrell Jr, 2007; Elbaum and Zondlo, 2014; Pandey *et al*., 2023). Finally, T259D and T259E, which were included as mimetics of the phosphorylated state of T259, also showed strongly impaired NPQ induction. Taken together, the strong impairment of NPQ induction in all T259 mutants for the first time demonstrates the importance of a stromal residue for the function of PsbS.

### PsbS C-terminus domain could be involved in hydrophobic protein-protein interactions

While all mutated T259 lines showed significantly slower NPQ induction, there were more nuanced effects observable associated with specific mutations, which may offer clues to unravel the role of the C-terminus in PsbS capacity to trigger NPQ. In particular, induction of NPQ in T259S and T259A was significantly less impacted than T259D and T259E, which showed the strongest impairment. Aspartate and glutamate have significantly lower side-chain hydrophobicity indices at pH of 7 than serine, alanine and threonine (Monera *et al*., 1995). Phosphorylation of threonine also significantly reduces side-chain hydrophobicity due to the addition of the charged phospho group. In all cases, these local effects may both increase or decrease hydrophobicity at the protein level, depending on structural rearrangements. However, since T259 in PsbS is followed by five adjacent hydrophilic glutamate and aspartate residues immediately downstream, it seems likely that a decrease in hydrophobicity at T259 via amino acid substitution or phosphorylation, would also decrease the hydrophobicity of the whole C-terminus.

Interestingly, molecular dynamics simulations of the impact of phosphorylation on a range of contrasting proteins showed that phosphorylated structures often exhibited reduced molecular flexibility, potentially due to additional contacts between protein residues invoked by the high charge of the phospho-group (Polyansky and Zagrovic, 2012), which may negatively affect interactions with surrounding proteins. Indeed, substitution of threonine with more hydrophilic residues has been observed previously to disrupt hydrophobic interactions of human *Heat Shock Protein 90* (Hsp90) with its co-chaperones (Mollapour *et al*., 2012) in a manner that mirrors our results. Similar to our findings, neither alanine or glutamate substitutions of threonine-22 rescued fully competent Hsp90, leading the authors to propose that proper conformation would require dynamic phosphorylation and dephosphorylation. If our observations with regards to T259 on PsbS are interpreted similarly, one could speculate that the C-terminus stromal domain of PsbS is involved in hydrophobic protein-protein interactions affecting induction of NPQ, which may be impacted by residue change at or phosphorylation of T259. Alternatively, the proximity of T259 to the β2 strand directly upstream, may interfere with interactions between β1 and β2, which in turn may impact protein-protein interactions.

In either case, the next logical question is which proteins could PsbS be interacting with to explain the effects on NPQ induction and recovery? One possibility is the propensity of PsbS dimers to undergo monomerisation (**Fig. 6A**), which would fit with the reduced NPQ induction phenotypes, considering the hypothetical role of dimer-to-monomer transition in qE formation (Bergantino *et al*., 2003; Pawlak *et al*., 2020). Alternatively, cross-linking and immuno-purification studies (Gerotto *et al*., 2015; Correa-Galvis *et al*., 2016 a; Sacharz *et al*., 2017) consistently show that PsbS promiscuously interacts with a range of LHC family proteins in darkness. While affinity for LHCB proteins generally increases when NPQ is induced (Correa-Galvis *et al*., 2016 a; Sacharz *et al*., 2017), LHCII M-trimers and the monomeric antenna LHCB4 (CP29) seem to show most evidence of reversible interactions upon return from light to dark conditions (Sacharz *et al*., 2017), in line with the reversible nature of qE. Indeed, when interactions were made irreversible by adding the hydrophobic cross-linker dithiobis(succinimidyl propionate) (DSP) to TM extracts, NPQ induction still occurred unperturbed, but recovery was completely abolished. Similarly, when the lumen-positioned H3 domain was removed (Chen *et al*., 2025), PsbS was still able to induce NPQ but recovery was significantly slower, implying that conformation changes at this domain (Liguori *et al*., 2019; Krishnan-Schmieden *et al*., 2021) may function to aid NPQ recovery by destabilising interactions between PsbS and LHCs upon de-protonation of the key glutamate-226. If the stability of PsbS-LHC interactions negatively impacts the rate of recovery of NPQ, the acceleration of NPQ recovery observed in substitution lines of T259 could reflect a destabilised interaction, leading to more rapid dissociation between PsbS and interacting LHC partner(s) when lumen pH returns to neutral (**Fig. 6B**). Interestingly, substitution of three phenylalanine residues for tyrosine in TM4 directly upstream from T259 led to strongly diminished NPQ induction (Chen *et al*., 2025), which was suggested to result from hampered hydrophobic interactions with LHCII trimers due to the increased polarity of the tyrosine residues. This would be consistent with a mechanism whereby substitution of T259 could hinder docking of LHCIIs on these protruding phenylalanines or interfere with stabilisation of the resulting complex.

**Fig. 6.**
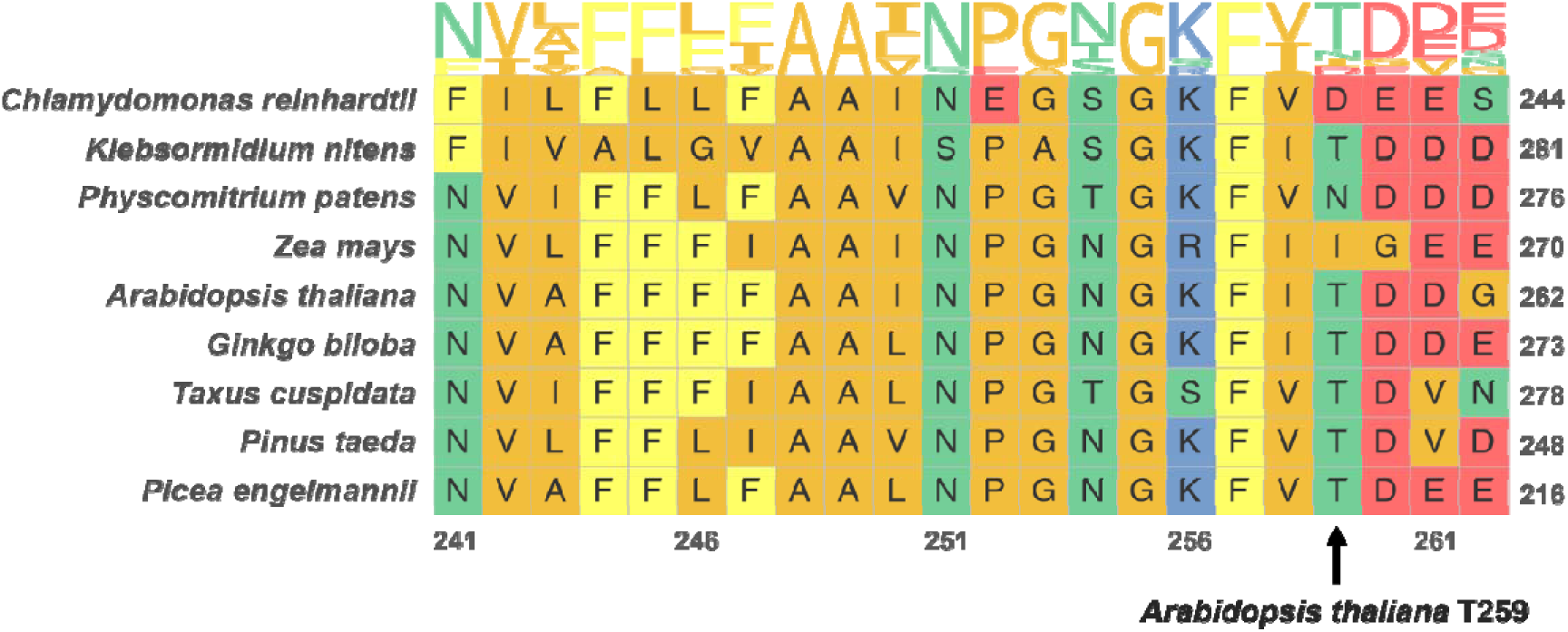
C-terminal PsbS sequence alignment of selected species from chlorophyte algae, streptophyte algae, bryophytes, gymnosperms, and angiosperms. The conserved residue (threonine-259 in *Arabidopsis thaliana*) is outlined in black. Sequence logo above the aligned sequences shows a visual representation of the proportion of certain amino acids at each site. Amino acids are coloured to reflect side-chain chemistry: negatively charged (red), positively charged (blue), non-polar aliphatic (orange), non-polar aromatic (yellow), and polar uncharged (green). Numbering on the x-axis is based on *A. thaliana* sequence, whereas on the y-axis, the corresponding sequence of individual species is denoted.

### De-activation of PsbS via reversible phosphorylation may be a specific feature of temperate evergreen conifers

T259 was first identified as a phosphorylation site by Roitinger *et al*. (2015) who performed a broad screen of phosphopeptides in Arabidopsis in response to ionising irradiance in wildtype and knockout mutants of the serine/threonine protein kinases *Ataxia Telangiectasia-Mutated* (*ATM*) and *Ataxia Telangiectasia-Mutated and Rad3-related* (*ATR*). While the C-terminal phosphopeptide containing T259 was identified in this study, there was no discernible effect of the radiation treatment and detection levels were also similar in the mutant lines compared to wild-type. Thus, neither condition provided evidence of reversible phosphorylation. Indeed, thus far the reversible phosphorylation of PsbS has only been reported for Norway spruce, an evergreen conifer adapted to survive the harsh winter conditions of extreme Northern latitudes. To survive these freezing conditions under illumination, these species appear to have evolved the ability to induce a sustained NPQ condition (quist and Huner, 2003). It seems plausible that under these circumstances, the facilitation of qE by PsbS prevents adequate acclimation. If so, phosphorylation of PsbS may provide a regulatory mechanism for the plant to toggle the qE effect of PsbS in harsh environments without degrading the protein, perhaps even helping to slow down the degradation of PsbS. Indeed, Merry *et al*. (2017) showed that certain light-harvesting proteins including PsbS were enriched in both white pine (*Pinus strobus L.*) and white spruce (*Picea glauca* (Moench) Voss), particularly during winter, which appears contrasting with the role of PsbS in qE, unless this pool is reversibly inactivated via phosphorylation, allowing more rapid recovery in spring without need for protein resynthesis.

While the threonine at T259 is conserved amongst the conifer species shown in **Fig. 7**, there are some notable different residues in this position across sequences from other selected species. The sequence from *Zea mays* (maize) carries an isoleucine instead of threonine and is flanked by a glycine, instead of aspartate. This could reflect further fine-tuning of PsbS action in species with C4 photosynthesis, but considering that the average hydrophobicity changes of both swaps combined would be minimal, it could also suggest that the overall hydrophobicity and associated conformation of the whole C-terminus is the leading factor, rather than the specific residue at 259. Interestingly, while the sequence from the streptophyte *Klebsormidium nitens* shows complete conservation of the C-terminus compared with Arabidopsis, the sequence from the model chlorophyte *Chlamydomonas reinhardtii* carries an aspartate at T259. Based on our results, the latter would strongly impair its ability to facilitate NPQ induction, which seems in line with experimental findings (Correa-Galvis *et al*., 2016 b; Redekop *et al*., 2020). Finally, the sequence from the model moss species *Physcomitrium patens* contains an asparagine instead of threonine at position 259, as well as valine instead of isoleucine at 258. While neither may have strong structural implications, it is also worth noting that in previous work by Gerotto *et al*. (2015) proper induction of qE was still observed in transgenic lines in which the final four amino acids (261-264) were replaced with a 6x repeat His-tag. Since this replacement involved swapping four hydrophilic amino acids (DEEE) with 6 hydrophilic histidines, it can be seen as further support for the importance of hydrophobicity across the C-terminus.

**Fig. 7.**
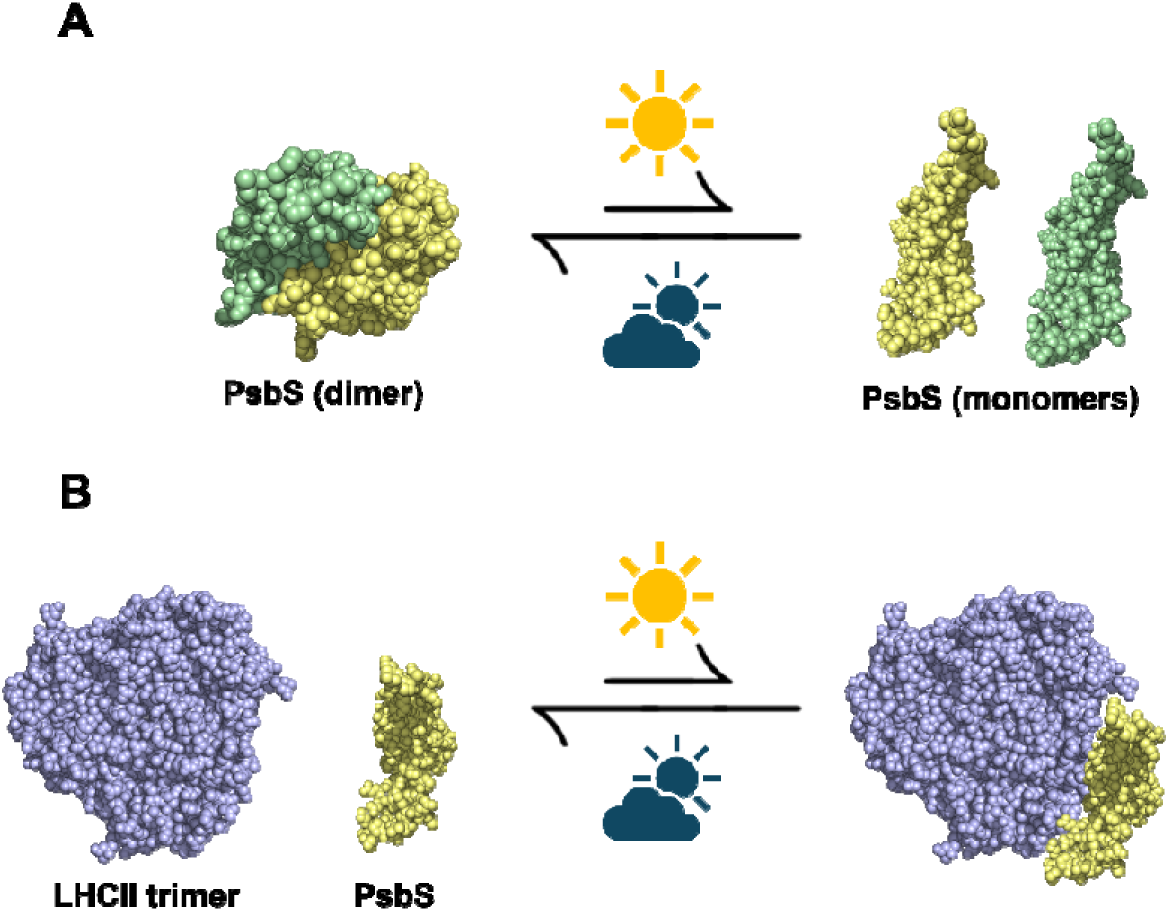
Proposed working model of T259-modified PsbS in downregulating NPQ. (**A)** Under high light and lumen acidification, T259-modified PsbS may disrupt the normal dimer-monomerisation kinetics, resulting in delayed activation of energy-dependent quenching (qE). **(B)** During qE formation, which involves architectural reorganisation of PSII-LHCII supercomplex essential for heat dissipation, T259-modified PsbS may exhibit reduced interaction efficiency with its LHC partners. Upon relaxation, T259-modified PsbS may dissociate from its LHC partners and revert to the dimeric form more rapidly than wild-type PsbS, explaining the generally faster relaxation observed in the T259 mutants. All illustrations are shown as top-view sphere representations. Protein models were built based on the following: PsbS dimer-PDB: 4RI2; PsbS monomer-UniProt: Q9XF91 (without transit peptide), LHCII trimer-PDB: 8YEE, and predicted using AlphaFold 3 (Abramson *et al*., 2024).

## Concluding remarks

In order to protect adequately against sharp light fluctuations or excess light, plants activate energy dissipation in the PSII antennae, which is facilitated by PsbS. While the mechanism of NPQ induction by PsbS involves protonation of key residues, here we followed up on the putative regulation via phosphorylation at T259, using a set of phosphomimetic lines. The strong decrease in NPQ induction and significant acceleration of NPQ recovery demonstrate for the first time the impact of the stromal-exposed C-terminus of PsbS. These results are consistent with hydrophobic interactions between PsbS and other LHC proteins possibly involving the fourth trans-membrane helix, which together with recent work elucidating key conformational changes associated with lumen-facing residues are finally starting to shape a mechanism underlying the action of this enigmatic protein.

## Supplementary data

**Table S1.** Lists of PsbS mutants, corresponding mutations, and the predicted functional effects.

**Table S2.** Lists of primers for the generation of PsbS mutants.

**Table S3.** List of primers used for the RT-qPCR analysis.

**Table S4.** ANOVA test results for parameters discussed in the main manuscript

**Fig. S1.** Sanger sequencing results for the validation of PsbS mutants.

**Fig. S2.** The maximum quantum yield of PSII after dark acclimation (Fv/Fm)

**Fig. S3.** The uncropped western blot membranes of total protein extracts

**Fig. S4.** The uncropped western blot membranes of thylakoid membrane extracts

## Author contributions

JK designed experiments, WYC performed all experiments and data analysis with help from JW and AKJR, WYC and JK drafted the manuscript, all authors contributed and approved the manuscript.

## Conflict of interest

None

## Funding

JK acknowledges a new lecturer start up award from the Gatsby foundation.

## Data availability

All data has been provided in the manuscript or supplemental materials.

## Notes

### Competing Interest Statement

The authors have declared no competing interest.

